# Structural basis for the binding of SNAREs to the multisubunit tethering complex Dsl1

**DOI:** 10.1101/2020.04.07.029496

**Authors:** Sophie M. Travis, Kevin DAmico, I-Mei Yu, Safraz Hamid, Gabriel Ramirez-Arellano, Philip D. Jeffrey, Frederick M. Hughson

**Author notes:** Corresponding author: Frederick Hughson.

## Abstract

Multisubunit tethering complexes (MTCs) are large (250 to >750 kDa), conserved macromolecular machines that are essential for SNARE-mediated membrane fusion in all eukaryotes. MTCs are thought to function as organizers of membrane trafficking, mediating the initial, long-range interaction between a vesicle and its target membrane and promoting the formation of membrane-bridging SNARE complexes. Previously, we reported the structure of the Dsl1 complex, the simplest known MTC, which is essential for COPI-mediated transport from the Golgi to the endoplasmic reticulum (ER). This structure suggested how the Dsl1 complex might function to tether a vesicle to its target membrane by binding at one end to the COPI coat and at the other end to ER SNAREs. Here, we use x-ray crystallography to investigate these Dsl1-SNARE interactions in greater detail. The Dsl1 complex comprises three subunits that together form a two-legged structure with a central hinge. Our results show that distal regions of each leg bind N-terminal Habc domains of the ER SNAREs Sec20 (a Qb-SNARE) and Use1 (a Qc-SNARE). The observed binding modes appear to anchor the Dsl1 complex to the ER target membrane while simultaneously ensuring that both SNAREs are in open conformations with their SNARE motifs available for assembly. The proximity of the two SNARE motifs, and therefore their ability to enter the same SNARE complex, depends on the relative orientation of the two Dsl1 legs.

## Introduction

Eukaryotic cells use vesicles to transport cargo between organelles and to the plasma membrane for exocytosis. These transport vesicles bear tail-anchored SNARE proteins which, in concert with complementary SNAREs in the target membrane, draw the two membranes into close apposition and facilitate membrane fusion. Each SNARE contains at least one SNARE motif, and sequence features within these motifs define four groups of SNAREs: Qa, Qb, Qc, and R (1,2). Fusogenic SNARE complexes contain one SNARE motif of each type and form via the coupled folding and assembly of cognate SNARE motifs into parallel 4-helical bundles. In addition to C-terminal transmembrane anchors and adjacent SNARE motifs, most SNAREs contain N-terminal regions that can regulate SNARE assembly and/or interact with other proteins. Many R-SNAREs contain N-terminal longin domains (3), whereas many Q-SNAREs contain N-terminal 3-helix bundles known as Habc domains (4-7). Some Qa-SNAREs exhibit “closed” conformations in which the Habc domain folds back onto the SNARE motif and prevents it from entering into a SNARE complex (8-11). Habc domains are also common within the N-terminal regions of Qb- and Qc-SNAREs (6,7,12,13), where the available evidence suggests that they function as protein-protein interaction modules (7,14).

In vivo, SNAREs collaborate with other factors including Sec1/Munc18 (SM) proteins and multisubunit tethering complexes (MTCs). SM proteins function as SNARE chaperones, regulating the assembly of SNARE complexes (11,15-17), whereas MTCs act upstream of SNARE complex assembly, mediating the initial attachment of a vesicle to its target membrane (17-19). Vesicle tethering canonically involves interactions between MTCs and membrane-associated Rab proteins, but can also involve MTC•SNARE, MTC•coat, MTC•golgin, and/or MTC•phospholipid interactions. MTCs also appear to participate directly in SNARE complex assembly and in formation of the fusion pore (20-24). As a step toward understanding these roles in greater detail, we focus here on structural studies of MTC•SNARE complexes. Only a few such structures have previously been reported, and in no case has it been possible to integrate them into a complete structure of the MTC (6,12,22,25).

The largest family of MTCs is the CATCHR (Complexes Associated with Tethering Containing Helical Rods) family, whose members function largely in trafficking to and from the Golgi (19,26,27). The CATCHR-family complexes – GARP/EARP, exocyst, conserved oligomeric Golgi (COG), and Dsl1 – contain 3-8 subunits each. Many of these subunits share a similar tertiary structure, the so-called CATCHR fold, comprising a series of helical bundles (28). The CATCHR fold is also found in the synaptic vesicle priming protein Munc13 (29). With the exception of the exocyst complex, the CATCHR complexes are multi-legged and flexible (20,30,31). These attributes seem well-suited for roles in vesicle docking and SNARE complex assembly.

Of the five CATCHR-family MTCs, the Dsl1 complex is the smallest (∼250 kDa in *Saccharomyces cerevisiae*), the simplest (just 3 subunits), and the only one for which an essentially complete high-resolution structural model (based on overlapping crystal structures at resolutions ranging from 1.9 to 3.0 Å) is available (20,32). The central subunit, Dsl1, bridges the other two subunits, Tip20 and Sec39, which each form an elongated leg. These legs, because of a flexible hinge within the Dsl1 subunit, are able to adopt a broad range of relative orientations (20). A distal portion of the Tip20 leg binds to the ER Qb-SNARE Sec20, while a distal portion of the Sec39 leg binds to the ER Qc-SNARE Use1 (20,32). The three Dsl1 complex subunits and the two SNAREs form a stoichiometric complex in vitro and in vivo (20,33,34), consistent with the idea that the Dsl1 complex is stably anchored to the ER membrane via interactions of its two legs with two different ER-resident SNAREs. With its legs in a parallel orientation, the Dsl1 complex is about 20 nm tall. At the apex of the complex, near the hinge, is an intrinsically disordered segment of the Dsl1 subunit called the lasso. The lasso contains singleton- and di-tryptophan motifs that bind to the COPI subunits α- and δ-COP, respectively (35-39). Thus the Dsl1 complex, by interacting at its base with ER SNAREs and at its tip with COPI vesicles, may mediate vesicle capture upstream of SNARE complex formation.

The interactions between the Dsl1 lasso and COPI coat subunits have been relatively well-studied by methods including x-ray crystallography (38,39). Here, we have used structural and biochemical experiments to reveal the molecular nature of the interactions between the two legs of the Dsl1 complex and the ER SNAREs Sec20 and Use1. We find that each interaction involves a tri-helical region of the corresponding SNARE. The observed binding modes would prevent each SNARE from adopting the closed conformation that has been observed for Qa-SNAREs, thereby leaving its SNARE motif free to engage other SNAREs. Placed in the context of the intact Dsl1 complex, the structures we report are consistent with the previous observations that the Dsl1 complex accelerates SNARE complex assembly, albeit modestly, and can bind fully assembled SNARE complexes (20). More generally, our results highlight the critical roles of SNARE N-terminal domains in mediating interactions with other elements of the vesicle docking and fusion machinery.

## Results

### Tip20•Sec20_NTD_ structure

The *S. cerevisiae* Dsl1 complex subunit Tip20 was discovered as a cytoplasmic protein that interacts with the cytoplasmic domain of the ER Qb-SNARE Sec20 (40). Biochemical experiments established that Tip20 binds an N-terminal region (residues 1-175), but not the SNARE motif, of Sec20 (20). Conversely, the N-terminal region of Tip20 (residues 1-81), which mediates binding to Dsl1, was not needed for binding to Sec20 (32). Therefore, we initially conducted crystallization screens using both full-length *S. cerevisiae* Tip20 and an N-terminally truncated version in combination with various constructs representing the N-terminal region of Sec20. Although a number of crystals were obtained, none of them diffracted well enough to allow structure determination. As an alternative approach, we screened orthologous Tip20•Sec20 complexes from other yeasts, co-expressed in bacteria. For this screen we used full-length Tip20 and an N-terminal fragment of Sec20 terminating just before the SNARE motif (Sec20_NTD_). While most of the Tip20•Sec20_NTD_ complexes were stable and soluble, only the *Eremothecium gossypii* complex yielded crystals. Although these initial crystals diffracted poorly, replacing full-length Tip20 with an N-terminally-truncated variant (Tip20_A-E_; Figure 1*A*) led to the discovery of an additional crystal form that diffracted x-rays to 3.2 Å resolution.

**Figure 1.**
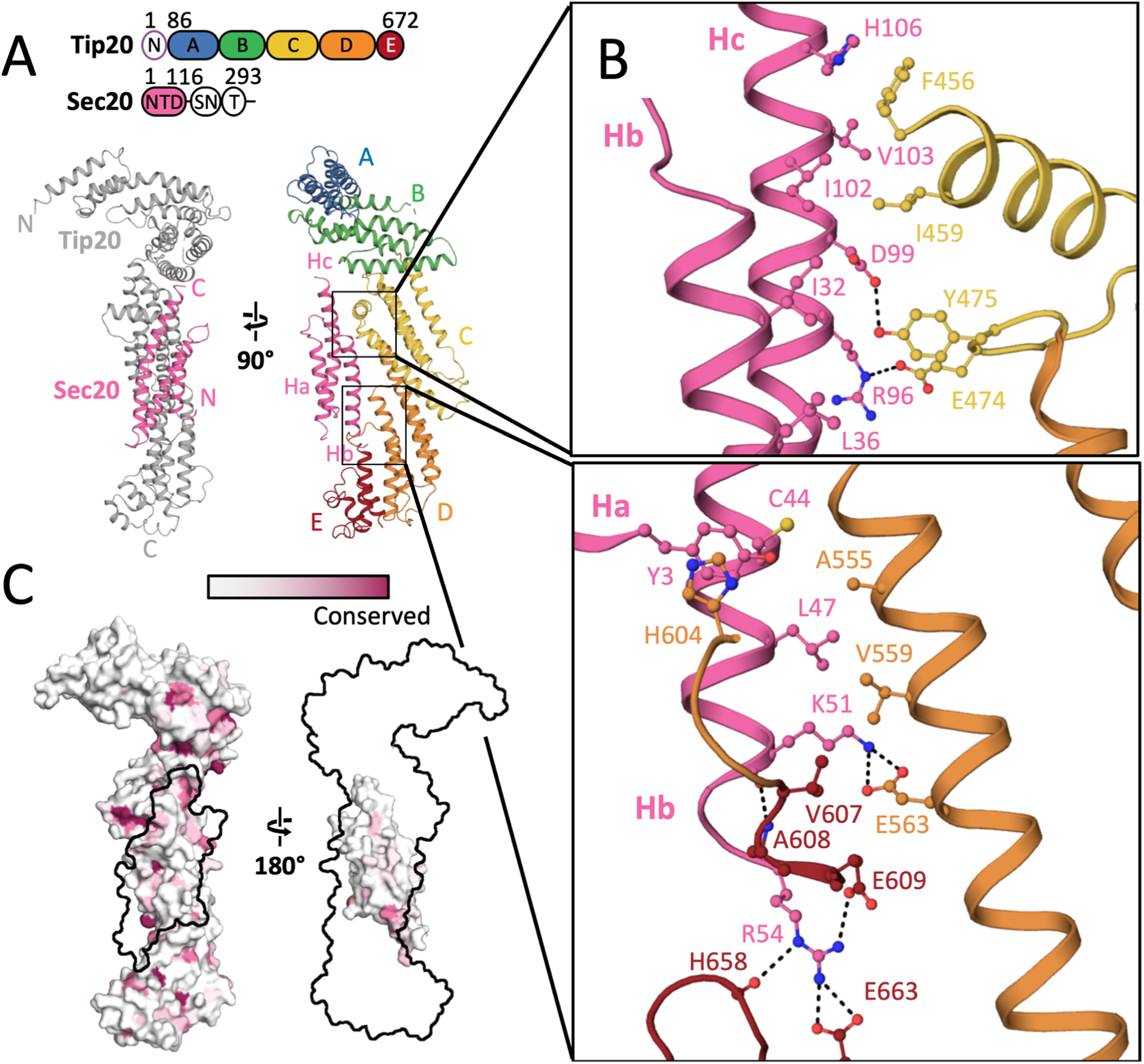
Structure of E. gossypii Tip20_A-E_•Sec20_NTD_ complex. *A*, Sec20_NTD_ adopts an Habc fold, interacting with Tip20 domains C through E. The full domain architecture of the two proteins is cartooned above. The Dsl1 complex subunit Tip20 consists of an N-terminal Dsl1-interacting domain (N), followed by 5 CATCHR domains (A-E). The SNARE protein Sec20 consists of an N-terminal Habc domain (NTD), SNARE motif (SN), and transmembrane helix (T). *B*, Details of the Tip20•Sec20 interface. *C*, The interacting surfaces of Tip20 and Sec20 are not highly conserved.

*E. gossypii* Tip20_A-E_•Sec20_NTD_ was phased by molecular replacement using the previously reported *S. cerevisiae* Tip20 structure as a search model (32). In the resulting structure (Figure 1*A*, Table S1), *E. gossypii* Tip20_A-E_ displays the characteristic hooked structure that appears to set Tip20 apart from other CATCHR-fold proteins (32) (Figure S1*A*). The greatest difference between *E. gossypii* Tip20 (in complex with Sec20_NTD_) and *S. cerevisiae* Tip20 (uncomplexed) lies in the angle between domains B and C (Figure S1*A*). Although this might reflect an inherent difference between the two orthologues, it could also reflect intrinsic flexibility at the B/C domain junction. A comparable degree of flexibility near the B/C junction was also observed for the exocyst subunit Exo70 (41), while dramatic flexing near the B/C junction of the Dsl1 subunit gives rise to the hinge motion that allows the entire Dsl1 complex to open and close (20).

Besides Tip20_A-E_, we observed electron density for three additional α-helices. Side chain density allowed these helices to be unambiguously assigned to Sec20_NTD_ (Figure S1*B*). The helices adopt a canonical Habc fold, with a disordered loop (residues 59-77) connecting Hb and Hc. Also disordered is the C-terminal region of the crystallized Sec20 fragment (residues 113 to 136), corresponding to the linker between the Habc domain and the SNARE motif. Previously, structures of Habc domains from several Qa-SNAREs, one Qb-SNARE, and one Qc-SNARE have been reported (4-7,12-14,42). Among these, the Habc domain of the Qb-SNARE Vti1 (13) bears the closest resemblance to that of Sec20, with a root mean squared deviation (rmsd) of 2.3 Å over 78 aligned C_α_ atoms (Figure S1*C*). This structural similarity hints that Habc domains may be a broadly conserved feature of Qb-SNAREs, as they are of Qa-SNAREs.

Sec20_NTD_ makes direct contact with Tip20 domains C-E, burying a surface of 1300 Å^2^ (Figure 1*A*). The interface is split into two parts, one a contact between domain C of Tip20 and helices Ha and Hb of Sec20_NTD_ and the other a contact between domains D and E of Tip20 and helices Hb and Hc of Sec20_NTD_ (Figure 1*B*). The centers of the two contact patches are formed by hydrophobic contacts between Ile-459 of Tip20 and Ile-102 of Sec20, and Ala-555 of Tip20 and Leu-47 of Sec20, respectively. These residues are comparatively well conserved, although Ala-555 is frequently replaced with a bulkier hydrophobic residue. The *E. gossypii* Tip20•Sec20 interface also features 6 salt bridges and 10 hydrogen bonds (Figure 1*B*).

### Tip20•Sec20_NTD_ binding

Unexpectedly, residues in the Tip20•Sec20_NTD_ binding interface are not especially well-conserved, compared to the Tip20 and Sec20 surfaces overall (Figure 1*C*), nor are they distinctive in terms of electrostatic surface potential or hydrophobicity (Figure S2*A*). Therefore, we sought to verify that the interface observed in our crystal structure is indeed responsible for complex formation in solution. The PISA server (43) predicted that the observed interface has a high probability of being biologically relevant and further identified three of the four residues cited above (Ile-459 in Tip20 and Leu-47 and Ile-102 in Sec20; Figure 1*B*) as among the residues contributing most favorably to the free energy of binding. We therefore substituted each of these residues, one at a time, with aspartate and used size exclusion chromatography to evaluate Tip20•Sec20_NTD_ complex formation. Binding of *E. gossypii* Tip20 to MBP-Sec20_NTD_ (maltose binding protein fused to Sec20_NTD_) was robust (Figure 2*A*). Each of the three substitutions, however, abolished binding (Figure 2*B-D*). These results demonstrate that the interface we observe is not a crystallization artifact. We also characterized binding of the wild-type partners by isothermal titration calorimetry (ITC) and found that the dissociation constant is approximately 100 nM (Table S2, Figure S2*B*).

**Figure 2.**
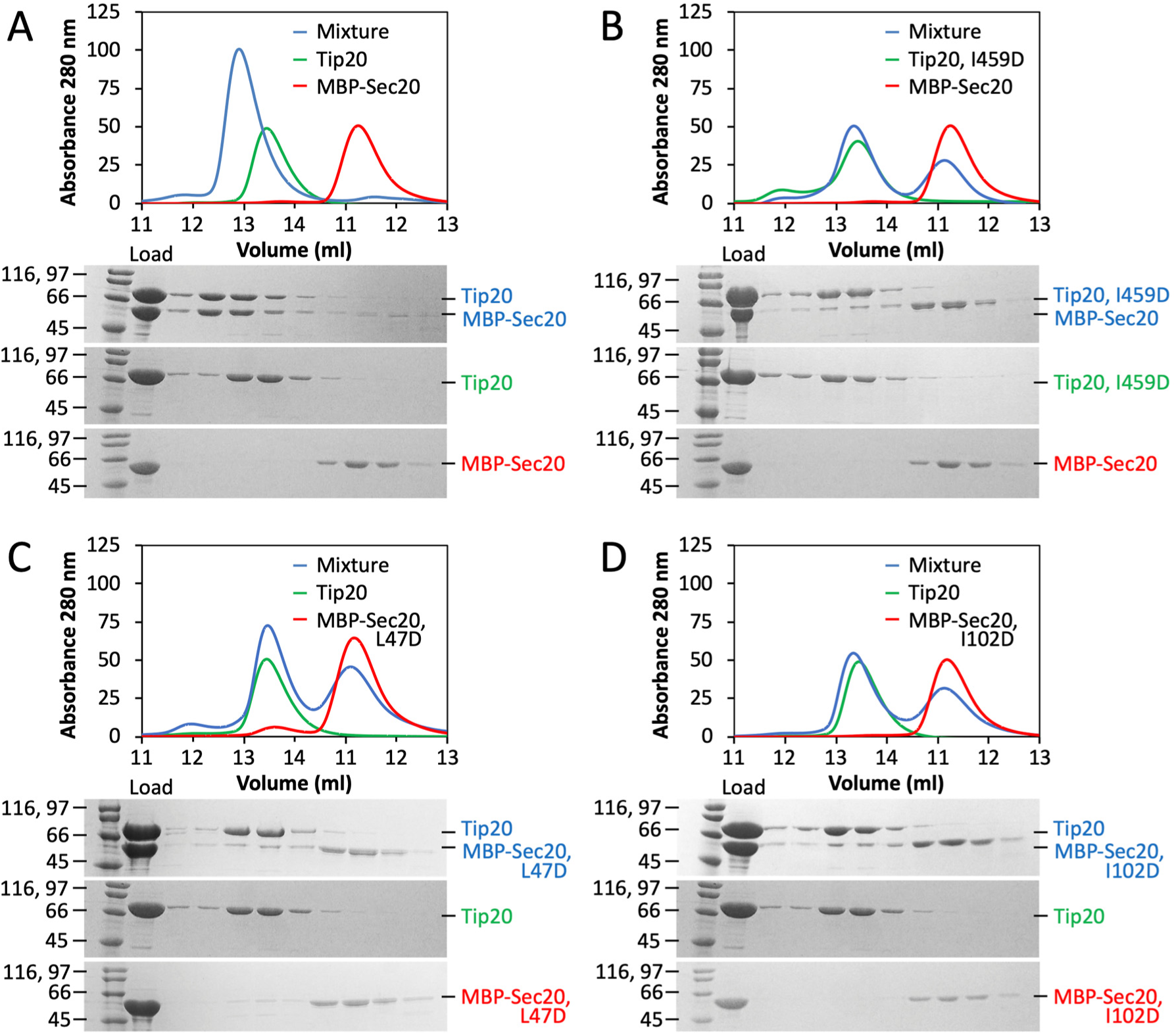
Structure-guided disruption of *E. gossypii* Tip20•Sec20 complex formation. *A*, Size exclusion chromatography demonstrates robust binding (blue) between wild-type *E. gossypii* Tip20 (green) and MBP-Sec20_NTD_ (red). *B-D*, Substitution of Tip20 Ile-459, Sec20 Leu-47, or Sec20 Ile-102 with aspartate abolishes binding of Tip20 to MBP-Sec20.

As noted above, Tip20 and Sec20_NTD_ orthologues from a broad sampling of yeast species form stable complexes, suggesting that the interface may be conserved despite the modest sequence conservation of the interfacial residues. To test this more directly, we conducted binding experiments using *S. cerevisiae* proteins. Structurally equivalent residues in *S. cerevisiae* and *E. gossypii* Tip20 were identified on the basis of their x-ray structures. Unexpectedly, replacing *S. cerevisiae* Tip20 residues Ile-481 and Leu-585 (equivalent to *E. gossypii* Ile-459 and Ala-555) with aspartate, individually or in combination, had little effect on Tip20•Sec20_NTD_ complex formation as judged by size exclusion chromatography (Figure S3). A more quantitative analysis using ITC revealed, however, that the double replacement led to a 15-fold increase in the equilibrium dissociation constant (Table S2, Figure S4). Predicting interface residues in *S. cerevisiae* Sec20_NTD_ was more challenging; not only is the sequence homology low, but we lack a *S. cerevisiae* Sec20_NTD_ structure to compare with *E. gossypii* Sec20_NTD_. Nonetheless, we found that changing *S. cerevisiae* Sec20 residue Val-82 (likely equivalent to *E. gossypii* Leu-47), alone or in combination with Leu-149 (likely equivalent to *E. gossypii* Ile-102), to aspartate modestly compromised Tip20•Sec20_NTD_ complex formation as judged by size exclusion chromatography (Figure 3). Overall, our data suggest that interface mutations designed on the basis of the *E. gossypii* x-ray structure destabilized the *S. cerevisiae* complex, implying that, despite the lack of strong sequence conservation, the interface is conserved. These results are suggestive of relatively rapid co-evolution of the interfacial residues.

**Figure 3.**
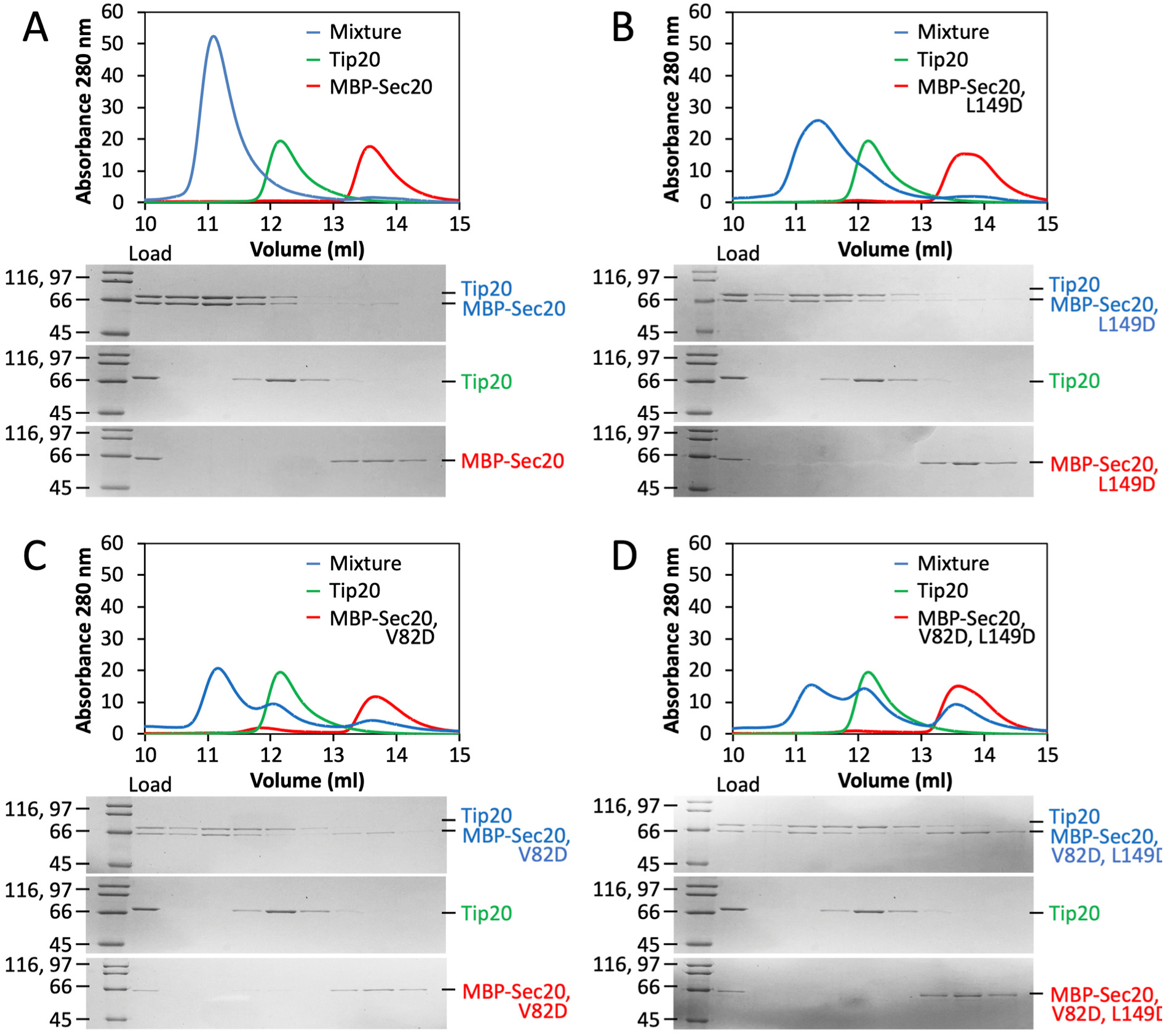
Structure-guided disruption of *S. cerevisiae* Tip20•Sec20 complex formation. *A*, Size exclusion chromatography demonstrates robust binding (blue) between wild-type *S. cerevisiae* Tip20 (green) and MBP-Sec20_NTD_ (red). *B*, Substitution of Sec20 Leu149 with aspartate has little effect on complex formation. *C*, Substitution of Sec20 Val-82 with aspartate partially compromises binding to Tip20. *D*, The two substitutions in combination have a stronger effect on binding.

To test the effect of weakening the Tip20•Sec20 interface in vivo, we used a plasmid shuffling strategy in *S. cerevisiae* to replace wild-type Tip20 or Sec20 with mutant versions. Deleting the N-terminal domain of Sec20 was lethal (Figure 4). Although this result is consistent with the model that the Tip20•Sec20_NTD_ interaction is essential in vivo, it is also possible that the Sec20 NTD plays an essential role that is independent of its binding to Tip20. Therefore, we attempted to confirm the model by replacing wild-type Tip20 or Sec20 with the double mutants characterized above. These substitutions did not, however, give rise to noticeable growth defects (Figures 4, S5). Thus, the affinity of Tip20 for Sec20 can be substantially reduced (ca. 15-fold for the structure-guided Tip20 mutations) without causing a major reduction in growth rate.

**Figure 4.**
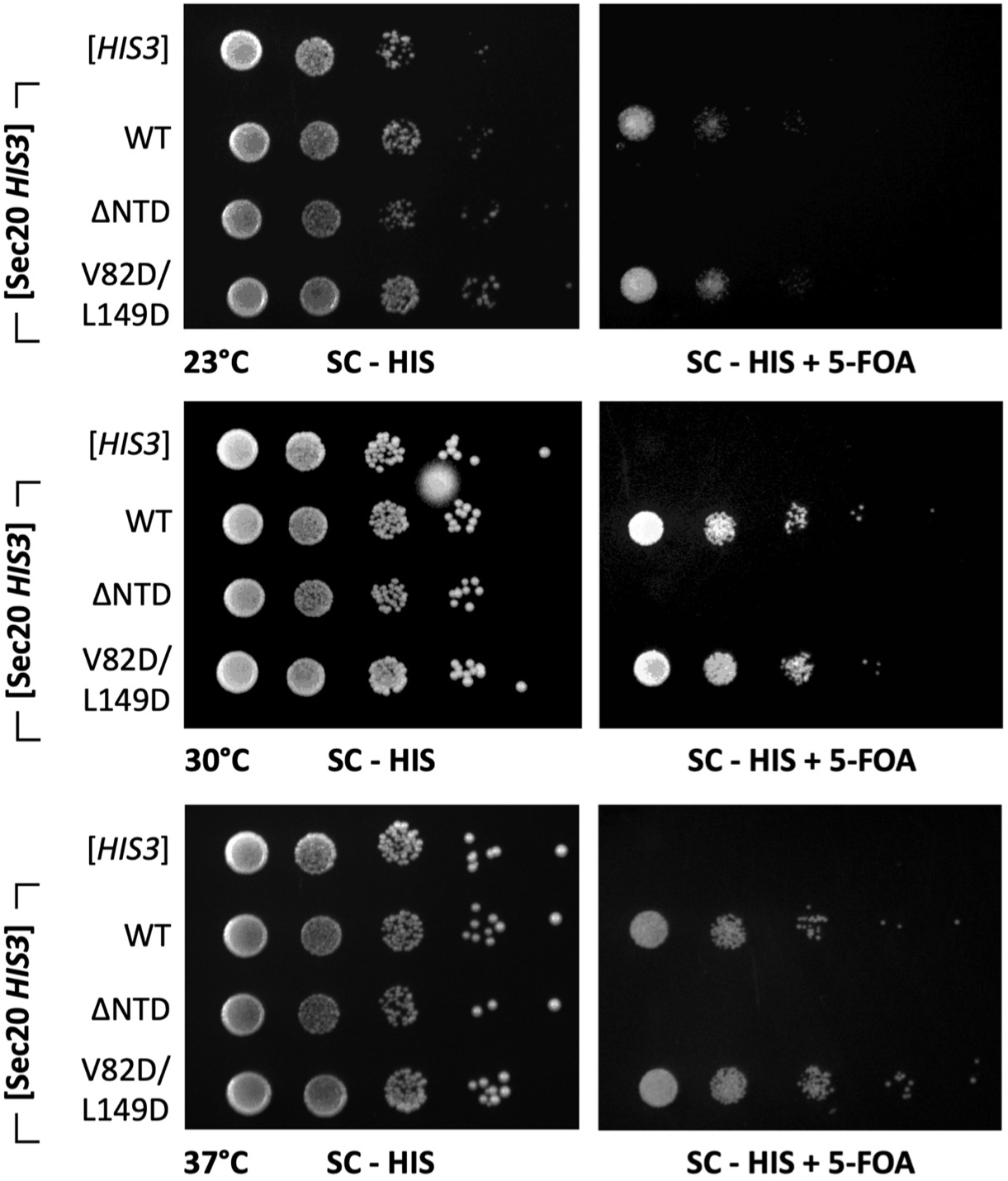
The N-terminal domain of Sec20 is essential for yeast viability. Yeast strains lacking endogenous Sec20 were maintained using a wild-type Sec20 covering plasmid marked with Ura3 and a second plasmid with the His3-linked Sec20 allele indicated at left. When the Ura3 plasmid is lost on 5-FOA selective plates (right), yeast harboring Sec20 lacking its N-terminal domain (ΔNTD) were inviable at all temperatures tested. Structure-based mutations in the Tip20•Sec20 interface (Tip20 V82D/L149D), which reduced the equilibrium dissociation constant by about 15-fold (see Table S2), did not have an evident impact on viability.

**Figure 5.**
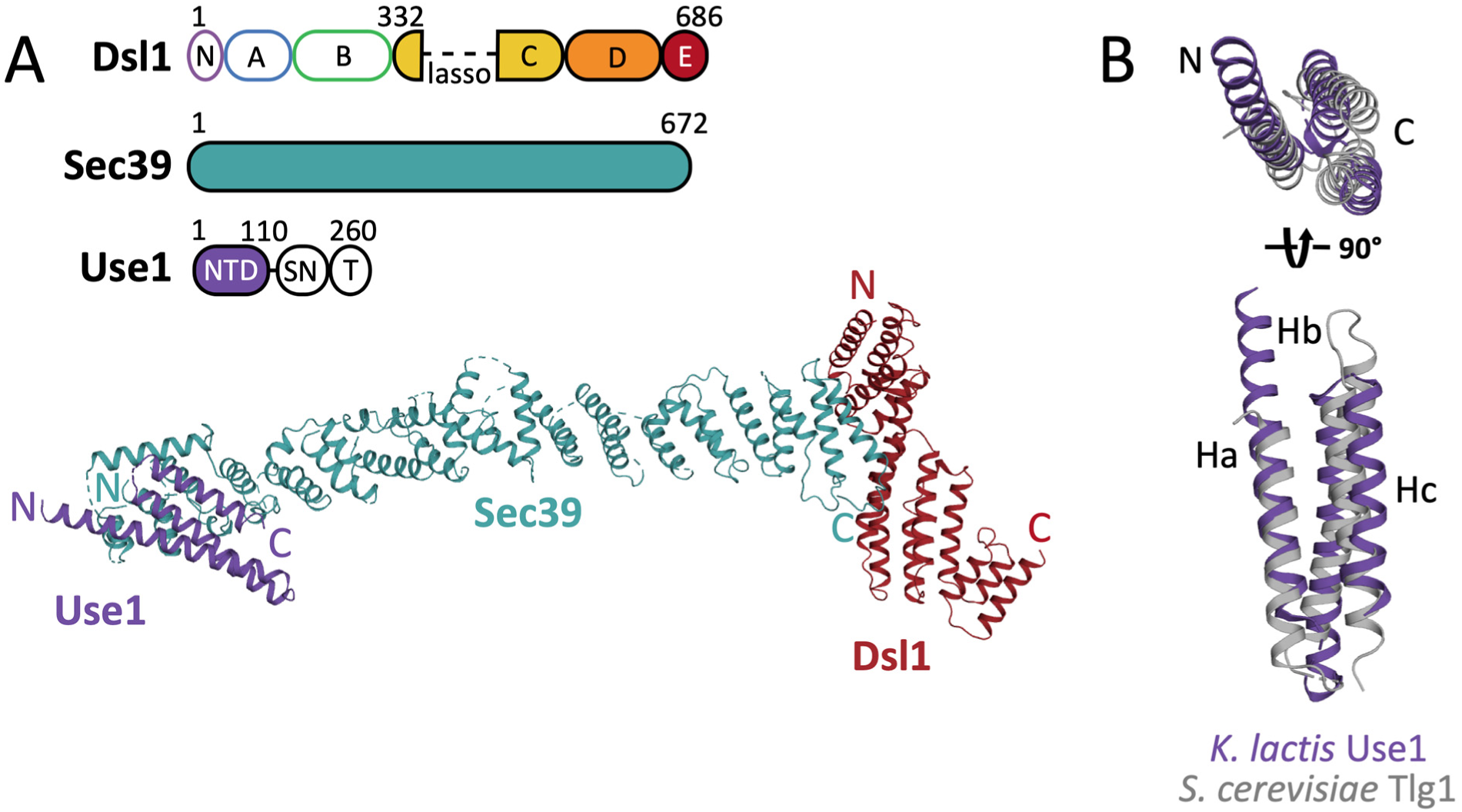
6.5 Å-resolution x-ray structure of *K. lactis* Sec39•Use1_NTD_•Dsl1_C-E_. *A*, The upper panel depicts the domain architecture of the three proteins crystallized. Dsl1 consists of an N-terminal Tip20-interacting domain (N) followed by 5 CATCHR domains (A-E). Sec39 consists of a long α-solenoid; for further details, see Figure S6. The SNARE protein Use1 consists of an N-terminal domain (NTD), a SNARE motif (SN), and a transmembrane helix (T). Below, Use1 (purple) binds to the extreme N-terminus of Sec39 (blue). **B**. K. *lactis* Use1 (purple) superimposes on its closest homologue, *S. cerevisiae* Tlg1 (gray, PDB accession code 2C5K, chain T) with an RMSD of 4.3 Å over 75 residues of the structure.

### Low-resolution Sec39•Dsl1_C-E_•Use1_NTD_ structure

The Sec39 leg of the Dsl1 complex is anchored to the ER membrane through an interaction with an N-terminal region of the Qc-SNARE Use1. Previous work (20) demonstrated that the first 35 residues of *S. cerevisiae* Use1 are required for binding to Sec39 in vitro. After extensive orthologue screening and construct optimization, we were able to generate crystals of a complex containing *Kluyveromyces lactis* Sec39, Use1 (residues 1-110), and Dsl1 (domains C-E, with the lasso deleted) that diffracted to 6.5 Å resolution. The diffraction data were post-processed using an anisotropic resolution cut-off (44), which included data extending to 5.9 Å resolution (Table S1).The *K. lactis* Sec39•Dsl1_C-_ E•Use1_NTD_ data were phased using molecular replacement based on our previous Sec39•Dsl1_C-E_ structure, which contained *S. cerevisiae* Sec39 (30% sequence identity with *K. lactis* Sec39) and *K. lactis* Dsl1_C-E_ (20). Despite the limited resolution of the data, a convincing molecular replacement solution was found in two steps, first placing the C-terminal portion of Sec39 and Dsl1_C-E_ and then placing the N-terminal portion of Sec39. In the resulting model, the N- and C-terminal portions of Sec39 form a continuous α-solenoid.

Our earlier Sec39•Dsl1_C-E_ structure lacked two regions near the N-terminus of Sec39, residues 1-29 and 63-100. Reexamination of the earlier electron density map, however, revealed weak density consistent with two additional helical hairpins. These previously-unmodeled hairpins were also visible in the Sec39•Use1_NTD_•Dsl1_C-E_ map (hairpins 1 and 3 in Figure S6). Because side chains cannot be seen at this resolution, the corresponding α-helices (α1, α2, α5, and α6), as well as an extension of α7, were modeled as polyalanine. Also visible in the Sec39•Dsl1_C-_ E•Use1_NTD_ map was clear electron density for three additional α-helices located adjacent to the N-terminal portion of Sec39 (Figure S7). This density has no counterpart in the earlier Sec39•Dsl1_C-E_ map, where its position is blocked by a crystal contact, and was consistent with the secondary structure prediction that *K. lactis* Use1_NTD_ is highly α-helical (Figure S8). The closest homolog of Use1 with a known structure is the Habc domain of *S. cerevisiae* Tlg1, another Qc-SNARE (6). Therefore, we attempted molecular replacement with various Habc domains and obtained the highest-scoring solution by using *S. cerevisiae* Vti1 as a search model (13) (Figure S7, Table S3). Because it is based on low-resolution data, the resulting model of the complete *K. lactis* Sec39•Dsl1_C-E_•Use1_NTD_ complex must be treated as preliminary. Nonetheless, it suggests that the interface between *K. lactis* Sec39 and Use1_NTD_ primarily involves helices α1-2 and α5-7 of Sec39 and Hb of Use1. Overall, our results unambiguously demonstrate that Use1 binds the distal tip of Sec39 using a tri-helical domain that is very likely to be an Habc domain.

## Discussion

The Tip20•Sec20_NTD_ and low-resolution Sec39•Dsl1_C-E_•Use1_NTD_ structures are, to our knowledge, the first structures of full-length MTC subunits bound to SNAREs. These and our previous structures (20,32) enabled us to construct a model for the complete Dsl1 complex anchored to the ER membrane by direct interactions between distal leg segments and ER SNAREs (Figure 6). The participating subunits – one (Tip20) a member of the CATCHR fold family, the other (Sec39) an α-solenoid – bind to N-terminal Habc domains of the Qb-SNARE (Sec20) and Qc-SNARE (Use1), respectively. In both the Tip20•Sec20_NTD_ and low-resolution Sec39•Dsl1_C-E_•Use1_NTD_ structures, the Hb/Hc groove of the SNARE NTD appears to be occluded by binding to the tethering complex subunit. It is not known whether uncomplexed Qb- and Qc-SNAREs adopt closed conformations analogous to those observed for Qa-SNAREs, in which the SNARE motif occupies the groove between the Hb and Hc helices of the Habc domain (9,10,22). However, in vivo the Dsl1 complex-anchoring SNAREs are almost certainly in open conformations, with their SNARE motifs accessible for SNARE assembly.

**Figure 6.**
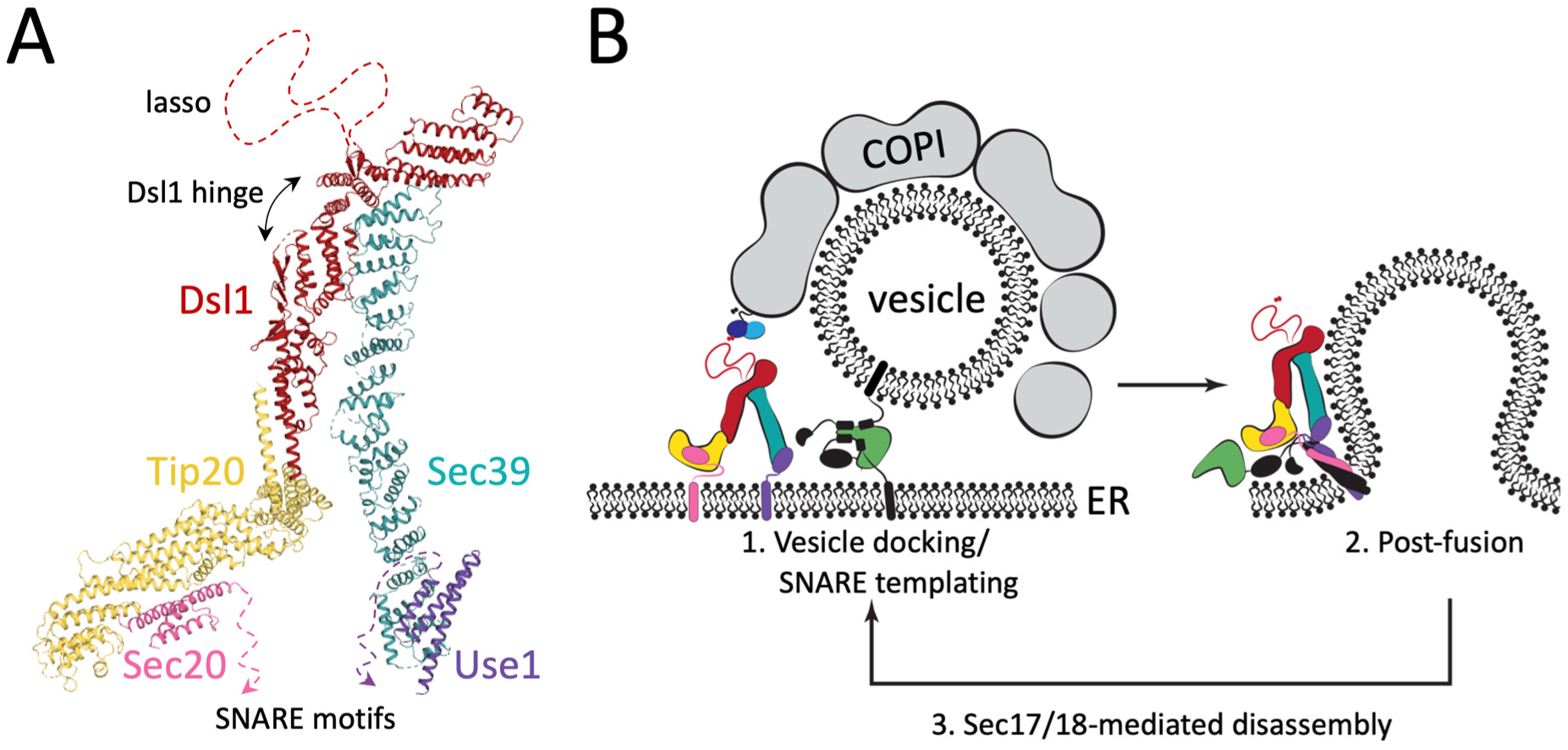
Proposed model for Dsl1-complex-mediated COPI vesicle tethering and fusion. *A*, A composite model of the complete Dsl1 complex in complex with the ER SNAREs Sec20 and Use1 was generated by combining the current structures with previously-reported structures of *S. cerevisiae* Dsl1 and Tip20 (PDB accession codes 3FHN and 3ETV). At the base of the complex the two SNARE motifs can extend towards one another and towards the ER membrane. At the top of the complex are the COPI-interacting lasso and the Dsl1 flexible hinge. *B*, The Dsl1 complex, colored as in (A), can adopt a range of conformations, positioning Sec20 and Use1 close enough to enter into the same SNARE bundle (right) or at a distance. Incoming vesicles are tethered by the Dsl1 lasso and, at later stages, a membrane-bridging complex is formed by the SM protein Sly1 (green) binding to SNAREs on the vesicle and ER membrane. The Dsl1 complex likely remains associated with the same SNAREs through multiple cycles of SNARE assembly and Sec17/18-mediated disassembly.

The Dsl1 subunit hinge angle would appear to be the most important variable determining whether the Sec20 and Use1 SNARE motifs are sufficiently close to allow assembly. Conformations in which the legs are roughly parallel will hold the SNARE motifs in relatively close proximity and would be expected to facilitate their entry into a SNARE complex together. On the other hand, with its legs in a sufficiently splayed conformation, the Dsl1 complex could prevent the bound Sec20 and Use1 SNAREs from participating in the same SNARE complex. An intriguing alternative is that the bound SNAREs might instead enter into two different SNARE complexes; multiple Dsl1 complexes might even generate an array (for example, a ring) of SNARE complexes. In any case, it is clear that factors that influence the Dsl1 hinge angle – or otherwise affect the distance between Sec20_NTD_ and Use1_NTD_ – could have a major effect on SNARE assembly.

Testing the physiological significance of the two tether-SNARE interactions was complicated by our inability to abolish them via the structure-guided design of interface mutations. In addition, the bivalent attachment of the Dsl1 complex to the ER membrane, via binding to two different ER SNAREs, may allow for vesicle tethering even when one of the anchoring interactions is compromised. Nonetheless, potential support for the importance of each Dsl1-SNARE interaction in vivo comes from the lethal effect of deleting the Sec20 Habc domain (Figure 4) and the strong temperature-sensitive growth defect caused by deleting the first 35 residues of Use1 (20). Both of these manipulations eliminate binding to the Dsl1 complex in vitro (20) but could in principle affect the localization and/or stability of the SNARE in vivo. Despite this limitation, our results highlight the functional importance of N-terminal domains not only for Qa-SNAREs (8,10,45), but for Qb- and Qc-SNAREs as well.

In their unfolded states, the Sec20 and Use1 SNARE motifs represent flexible connectors between the Dsl1 complex and the ER membrane. Anchoring each leg of the Dsl1 complex via a flexible segment would appear to extend the reach of the Dsl1 complex as a vesicle tethering factor. With its legs in a parallel orientation, the Dsl1 complex itself has a height of about 20 nm, and the disordered lasso at its tip extends this range further. The unfolded portion of the SNARE, at least 70 residues in length, would readily permit excursions of 10-15 nm from the ER membrane. Thus, the SNARE-anchored Dsl1 complex may be capable of capturing COPI vesicles that are 40 nm or more from the ER surface. Although this is a greater distance than SNAREs alone can reach (10-15 nm for unfolded SNARE motifs zippering from their N-terminal ends), it is far less than the tethering distance spanned by homodimeric coiled-coil tethering factors, which in fully extended conformations could reach 200 nm or more (46). Flexible regions within the Golgi coiled-coil tethering protein GCC185, however, allow its N- and C-termini to approach within about 40 nm of one another (47). Another well-studied coiled coil tethering protein, EEA1, undergoes entropic collapse upon vesicle binding (48), which could serve to pull the vesicle close enough to engage an MTC. In the case of the Dsl1 complex, retrograde trafficking from the Golgi to the ER has not been shown to require a coiled coil tethering protein. It is possible that the need for long-range tethering is obviated by the close association of ER export and arrival sites. In the yeast *Pichia pastoris*, this creates bidirectional transport portals at which the Dsl1 complex may be able to capture COPI vesicles as they are budding from the Golgi (49).

What of SNARE assembly itself? The Dsl1 complex forms a stoichiometric complex in vivo with Sec20 and Use1 (33). Analogous complexes (containing the cytoplasmic portions of the SNAREs) are stable in solution and, in the additional presence of the Qa-SNARE Ufe1 and the R-SNARE Sec22, assemble to form a heptameric complex containing all four SNAREs and all three Dsl1 subunits (20). Based on these results and our current model for the anchored Dsl1 complex, we suggest that SNAREs undergo cycles of assembly and Sec17/18-catalyzed disassembly while remaining associated with the Dsl1 complex. In this way the Dsl1 complex, by directing vesicles to sites in which at least two of the three Q-SNAREs are already present, may be able sequentially to facilitate both tethering and SNARE assembly. It would however fall to other factors, such as the SM protein Sly1, to protect the assembling SNARE complexes from the disassembly activity of Sec17/18. Indeed, our results hint at a possible division of labor between two classes of protein – MTCs and SM proteins – known to chaperone SNARE complex assembly. As described here, the Dsl1 complex binds to Qb- and Qc-SNAREs in such a manner that their SNARE motifs are available for assembly. SM proteins, on the other hand, bind Qa-SNAREs, and recent studies of several SM proteins suggests that they bind R-SNAREs at an adjacent site and thereby function as templates to initiate SNARE assembly (16,25). Thus it is attractive to speculate that, upon close approach of the vesicle, its R-SNARE Sec22 engages the Sly1-bound Qa-SNARE Ufe1, to which the Dsl1 complex then presents the SNARE motifs of Sec20 and Use1 for the formation of a membrane-bridging trans-SNARE complex and subsequent membrane fusion.

## Methods

### Protein expression and purification

A DNA fragment corresponding to *E. gossypii* Tip20 (residues 1-672) was amplified from genomic DNA by PCR and cloned directionally into the pQLinkH expression vector (50) using the BamHI/NotI restriction sites. Similarly, Sec20 (residues 1-136) was inserted into the pQLinkN vector. The two vectors were concatenated for co-expression using the LINK strategy (51). Subsequently, *E. gossypii* Tip20_A-E_ (residues 86-676) was subcloned into pQLinkH and concatenated with Sec20. The *K. lactis* expression vector was generated similarly, incorporating Sec39 into pQLinkH, Use1 (residues 1-110) into pQLinkN, and Dsl1 (residues 332-686, harboring an internal deletion of residues 367-423 replaced with the sequence Gly-Asp-Gly-Asp-Gly) into pQLinkN. The *S. cerevisiae* Tip20 and Sec20_1-174_ expression constructs have been previously described (20,32). For binding assays, Sec20 (residues 1-136) was subcloned into a modified pQLinkH expression vector containing an N-terminal MBP (25). All expression constructs were validated by DNA sequencing. Mutations were introduced using a modified QuikChange mutagenesis protocol (52).

All proteins were overproduced in BL21-CodonPlus(DE3) RIL *E. coli* (Agilent) grown in high salt Luria broth (Sigma) to an optical density at 600 nm (OD_600_) of 0.5-0.7 and induced by the addition of 0.5-1 mM isopropyl β-*D*-1-thiogalactopyranoside. Cells were harvested after 16 h induction at 15°C, resuspended in 20 mM HEPES pH 7.5, 150 mM NaCl (HBS) supplemented with 5 mM β-mercaptoethanol, 1 mM phenylmethylsulfonyl fluoride, and 10 μg/mL DNase I (Roche) and lysed using a cell disrupter. After clarification, target proteins were purified from cell lysate by Ni-iminodiacetic acid affinity chromatography (Clontech). However, *S. cerevisiae* Tip20 was purified using TALON affinity chromatography (Clontech) in 20 mM Tris pH 8.0, 200 mM NaCl (TBS). All proteins except *S. cerevisiae* Tip20 were further purified by anion exchange chromatography (MonoQ, GE Healthcare). Finally, proteins were purified by size exclusion chromatography (Superdex200 Increase, GE Healthcare). Purified proteins were concentrated, flash-frozen, and stored at −80°C in TBS (*S. cerevisiae* Tip20) or HBS (all other proteins) supplemented with 1 mM DTT.

### Crystallization and data collection

Crystals were obtained of the *E. gossypii* Tip20_86-_ 672•Sec20_1-136_ heterodimeric complex by hanging drop vapor diffusion at 20°C, mixing 1 μL of protein (10 mg/mL) and 1 μL of well buffer (100 mM sodium citrate pH 6.0, 725 mM ammonium sulfate, 1 mM DTT). After 3 days, rounded hexagonal crystals 100 × 100 × 100 μm were obtained and cryoprotected using a 1:1 mixture of formulation buffer to well buffer supplemented with 30% v/v glycerol before flash freezing in liquid nitrogen. Data were collected at the US National Synchrotron Light Source II (NSLS-II) AMX beamline.

Crystals were obtained of the *K. lactis* Sec39•Use1_1-110_•Dsl1_332-686,ΔL_ heterotrimeric complex by hanging drop vapor diffusion at 20°C, mixing 1 μL of protein (5 mg/mL) and 1 μL well buffer (100 mM Tris pH 7.5, 100 mM NaCl, 8% w/v PEG 3,350, 10% v/v ethylene glycol, 1 mM DTT). After 1-2 days, thin plates 200 × 200 × 20 μm were obtained and cryoprotected in a solution containing 50 mM Tris pH 7.5, 10 mM HEPES pH 7.5, 125 mM NaCl, 11% w/v PEG 3,350, 30% v/v ethylene glycol and flash frozen in liquid nitrogen. Data were collected at the NSLS-II FMX beamline.

### Structure determination and refinement

Tip20_A-E_•Sec20_NTD_ data were processed using XDS (53). Initial search models for molecular replacement were derived from the *S. cerevisiae* Tip20 structure, PDB code 3FHN (32), with non-conserved loops deleted and sidechains pruned by the program CCP4 program CHAINSAW. Molecular replacement was carried out in PHASER (54), searching first for domains C-E, then separately for domain B. Rounds of refinement and manual building were performed using PHENIX.refine (55) and COOT, respectively (56).

Sec39•Dsl1_C-E_•Use1_NTD_ data were processed in XDS, and an anisotropic resolution cut-off was applied (44). Initial search models for molecular replacement were derived from the structure of *S. cerevisiae* Sec39 bound to *K. lactis* Dsl1_332-686_, PDB code 3K8P (20), modified by the CHAINSAW program. Molecular replacement in PHASER was carried in out in two steps, first searching for *S. cerevisiae* Sec39_380-672_ bound to *K. lactis* Dsl1_332-686_, then searching for *S. cerevisiae* Sec39_1-379_. Sec39 was maintained as two chains, one encompassing α1 to α19 and the other encompassing α20 to α34. One round of bulk solvent scaling, rigid body, and TLS (translation libration screw) refinement was performed in PHENIX.refine, assigning each chain as a single rigid body and a single TLS group. Four additional helices of Sec39 were built (α1, α2, α5, and α6) into unbuilt density in the *K. lactis* map, with reference to unbuilt regions of *S. cerevisiae* map. In addition, helices were extended or trimmed to fit the map and real space rigid body refinement was performed on poorly fit hairpins in COOT.

After a second round of refinement in PHENIX.refine, the updated model and map were used as a starting point for a new molecular replacement search in Phaser for each of the deposited Habc domain structures. The best solution, as judged by expected log likelihood gain, was found using *S. cerevisiae* Vti1 (13), PDB code 3ONJ (Table S3). The Vti1 model was adapted in COOT by extending the N-terminal helix and trimming the C-terminal helix to better reflect the density present in the *K. lactis* map. Each helix was rigid-body fit separately using COOT. Sec39 side-chains were not modeled beyond Cα unless identical to *S. cerevisiae* Sec39, in which case the *S. cerevisiae* conformation was used. A final round of single-chain rigid body and TLS refinement was performed as above. To verify the position of Use1, an omit map was calculated omitting the entire Use1 chain.

### Gel filtration binding assays

Proteins were diluted to 20 μM in HBS (*E. gossypii* proteins) or 8 μM in TBS (*S. cerevisiae* proteins) supplemented with 1 mM DTT in a volume of 200 μL and incubated at 4°C for 1 h, then loaded onto a Superdex 200 Increase 10/300 column. Fractions were collected and analyzed by 12% Tris/Glycine SDS-PAGE.

### Isothermal titration calorimetry

Proteins were exchanged into HBS (*E. gossypii* proteins) or TBS (*S. cerevisiae* proteins) supplemented with 1 mM DTT using pre-equilibrated BioSpin-6 spin columns (BioRad) or, for Tip20_L585D_ and Tip20_I481D,L585D_, 3,500 Da molecular weight cut-off Slide-A-Lyzer Mini Dialysis units (Thermo Scientific). Samples were then back-diluted to the indicated concentration. Tip20 was loaded into the sample cell and Sec20 orthologues into the titration syringe. Experiments were performed using a MicroCal PEAQ-ITC (Malvern) and analyzed using the MicroCal PEAQ-ITC Analysis Software package (Malvern).

### Yeast methods

A diploid strain heterozygous for the deletion of Sec20, marked with KanMX, was purchased from the essential gene knockout collection (Dharmacon). A Sec20 covering plasmid, containing the coding sequence of Sec20 and 500 bases upstream, was created based on the pRS416 Ura3-containing plasmid (57). After transformation of the covering plasmid (58), diploids were sporulated in sporulation media containing 0.3% w/v potassium acetate and 0.02% w/v raffinose at 23°C for 7 days. Haploid spores bearing a chromosomal deletion of Sec20 covered by the Ura3 plasmid were screened based on prototrophic and antibiotic resistance markers. Plasmids containing mutant alleles of Sec20, marked with His3, were then transformed into this haploid strain. Transformants were grown overnight at 30°C in synthetic complete media lacking histidine, then plated on synthetic complete agar lacking histidine, supplemented with 0.1% w/v 5-fluoroorotic acid (GoldBio) as indicated. The Tip20 deletion strain with Ura3 covering plasmid was created in an analogous manner. Mutants of Tip20 were introduced on a His3 plasmid and analyzed via 5-fluoroorotic acid counterselection.

### Figures

Sequence alignments were generated using ClustalW (59) and rendered using JalView (60). Secondary structure prediction was performed in Jpred (61). Superimposition was performed using the CCP4 utility Superpose with secondary structure matching. Structures were rendered using PyMol (Schrodinger), with conservation scores imported from ConSurf (62) where applicable. Hydrophobicity was assigned using the Eisenberg hydrophobicity scale (63).

### Data availability

The structures presented within this work have been deposited in the Protein Data Bank (PDB) with the following accession codes: 6WC3 for Tip20_A-E_•Sec20_NTD_ and 6WC4 for Sec39•Dsl1_C-E_•Use1_NTD_. All other data are available within the article.

## Acknowledgements

We thank Angela Chan, Carly Geronimo, Venu Vandavasi, Virginia Zakian, and members of the Hughson laboratory past and present for valuable advice and discussions. The Princeton Biophysics and Macromolecular Crystallography core facilities provided essential assistance with isothermal titration calorimetry and x-ray crystallography, respectively. We gratefully acknowledge the assistance provided by the AMX and FMX beamline staff at NSLS-II.

## Funding and additional information

This work was supported by National Institutes of Health (NIH) grants T32GM007388 (SMT and KD), F31GM12676 (SMT), and R01GM071574 (FH). This research used the AMX and FMX beamlines of the National Synchrotron Light Source II, a United States Department of Energy (DOE) Office of Science User Facility operated for the DOE Office of Science by Brookhaven National Laboratory under Contract No. DE-SC0012704. The Life Science Biomedical Technology Research resource, which supports AMX and FMX, is primarily supported by the NIH, National Institute of General Medical Sciences (NIGMS) through a Biomedical Technology Research Resource P41 grant (P41GM111244), and by the DOE Office of Biological and Environmental Research (KP1605010).

## Conflict of Interest

The authors declare no conflicts of interest with respect to this manuscript.

## Supporting Information

**Figure S1.**
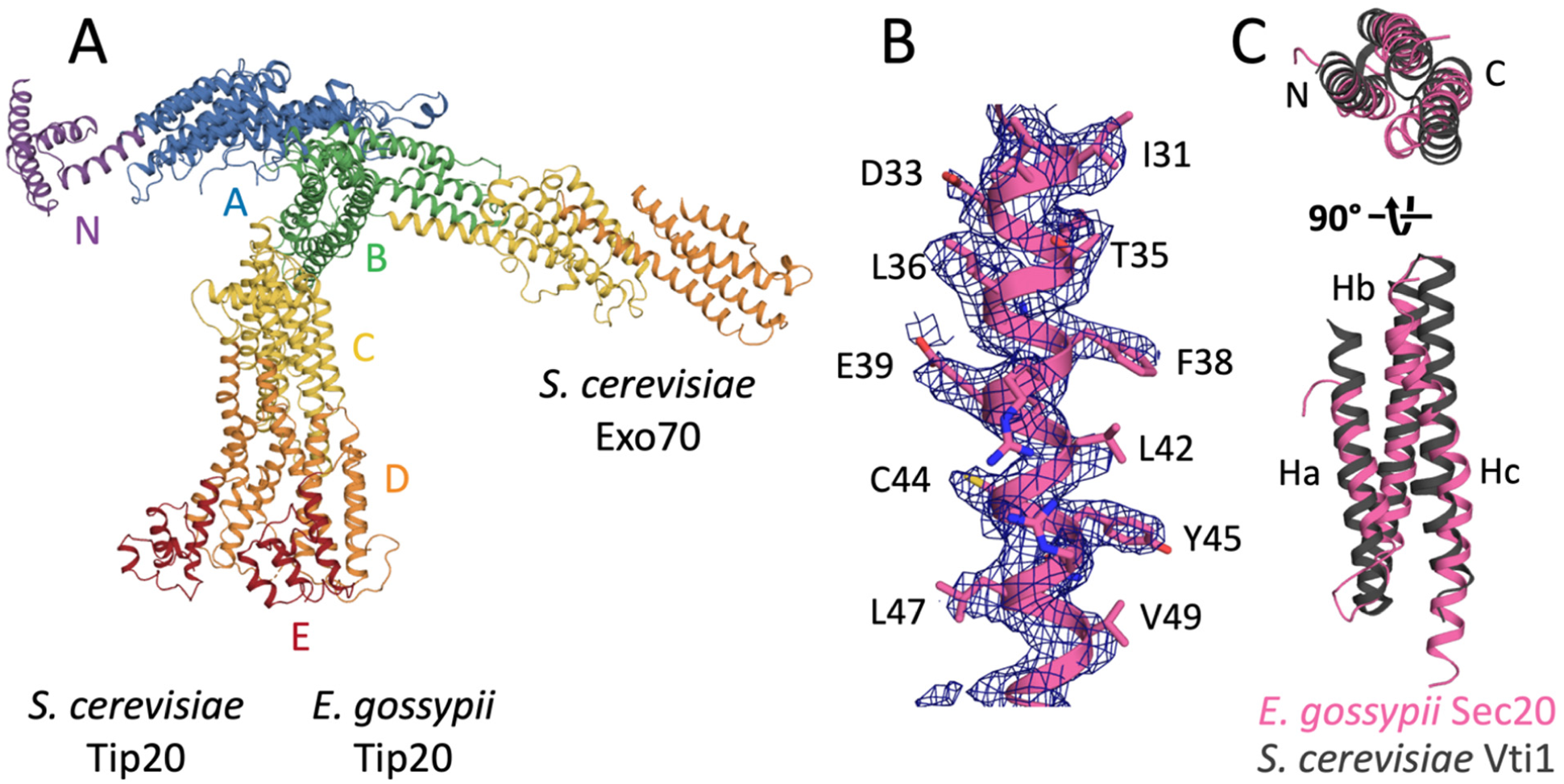
Comparative analysis of *E. gossypii* Tip20 and Sec20 structures. *A, E. gossypii* and *S. cerevisiae* Tip20 (3FHN, chain A) superimpose with an RMSD of 2.9 Å over 394 residues. The major difference between the two structures is the different relative orientation of domains B and C. *S. cerevisiae* Exo70 (2PFV) adopts a much more linear configuration. The three proteins are shown aligned on domain A. *B*, Representative electron density for *E. gossypii* Sec20 (pink), showing side chain assignments. The composite omit 2Fo-Fc map (dark blue mesh) is contoured at 1σ. *C, E. gossypii* Sec20_NTD_ (pink) resembles *S. cerevisiae* Vti1 (3ONJ, gray), superimposing with an RMSD of 2.3 Å over 78 residues of the structure.

**Figure S2.**
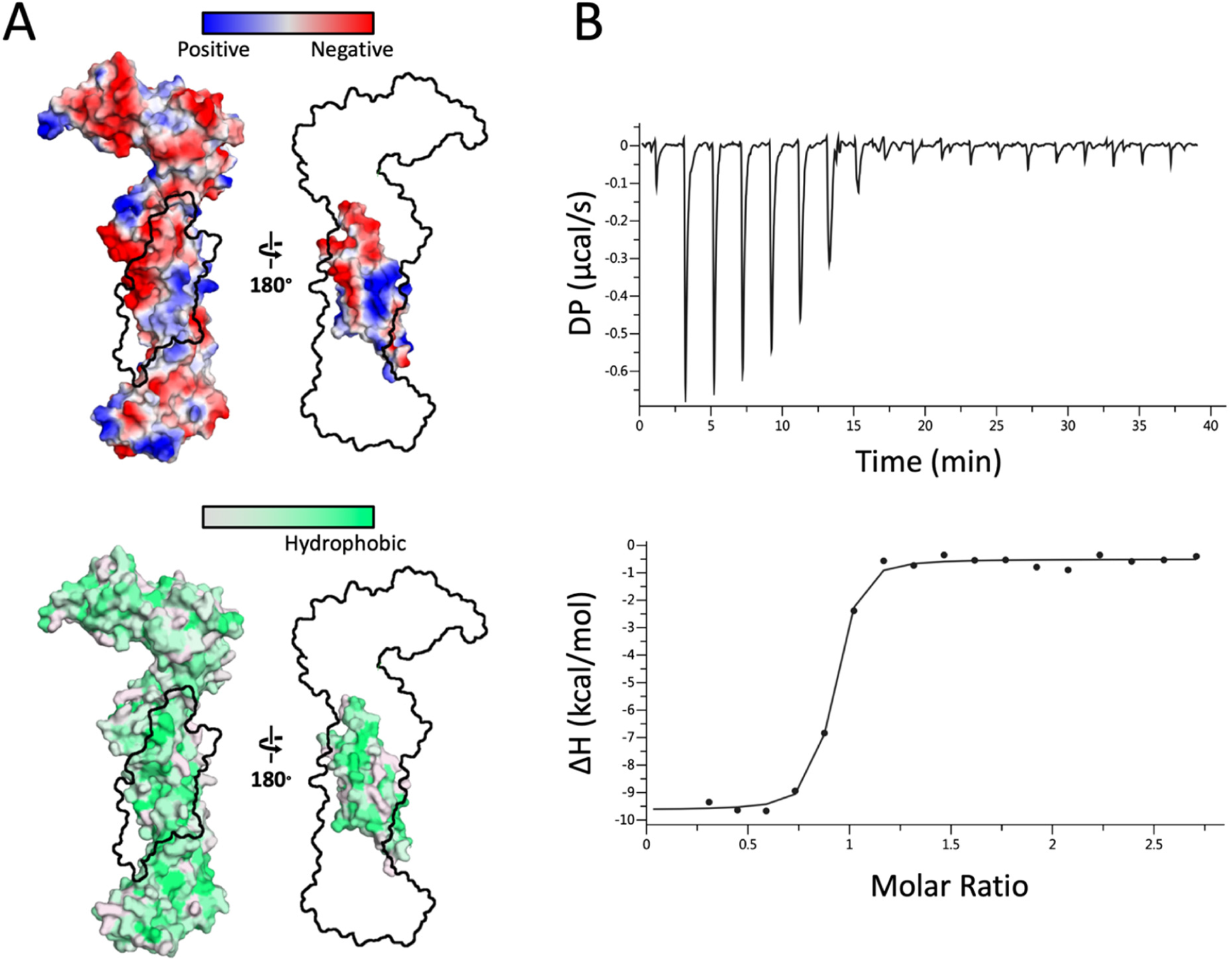
Properties of the *E. gossypii* Tip20•Sec20 complex. *A*, The protein:protein interface lacks obvious distinguishing features in terms of surface electrostatic potential (top) or hydrophobicity (bottom). *B*, Isothermal titration calorimetry data for MBP-Sec20_1-136_ and Tip20; see Table S2.

**Figure S3.**
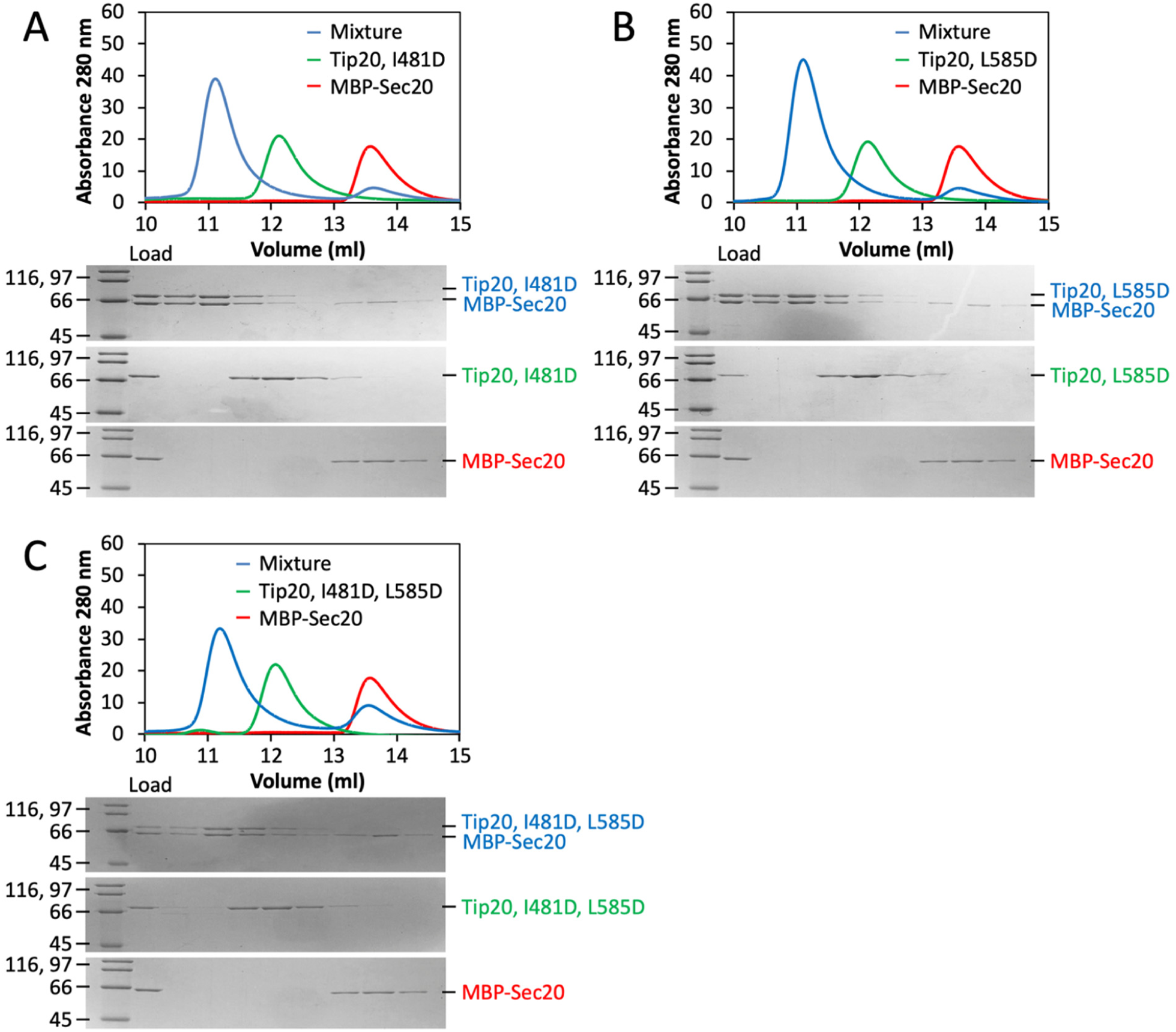
*S. cerevisiae* Tip20 mutants minimally affect binding as judged by size exclusion chromatography. *A-C*, Replacing *S. cerevisiae* Tip20 Ile-481, Leu-585, or both with aspartate had little effect on complex formation.

**Figure S4.**
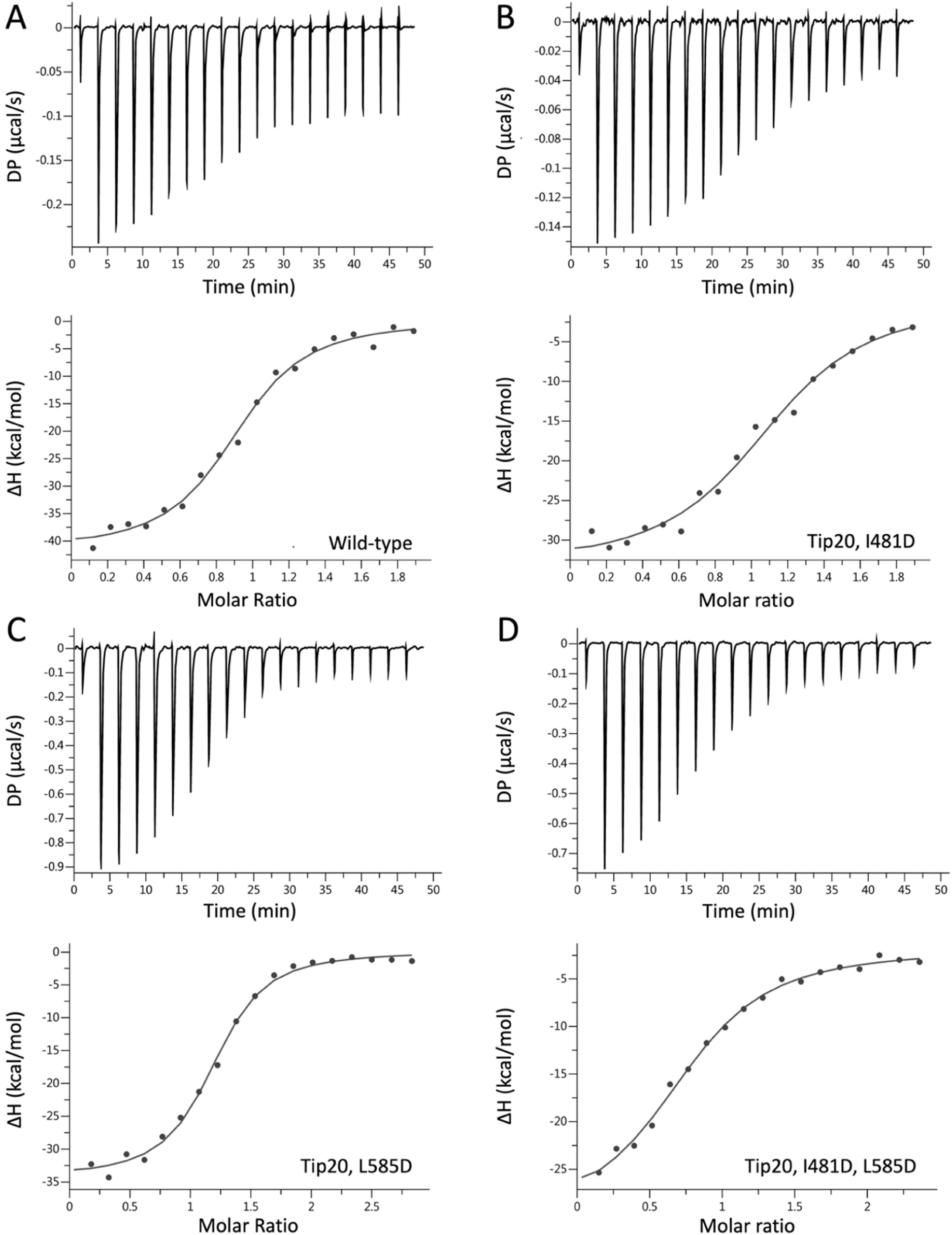
Structure-based mutations destabilize the *S. cerevisiae* Tip20•Sec20_1-174_ complex. *A-D*, Isothermal titration calorimetry data for Sec20_1-174_ and the indicated Tip20 mutants; see Table S2.

**Figure S5.**
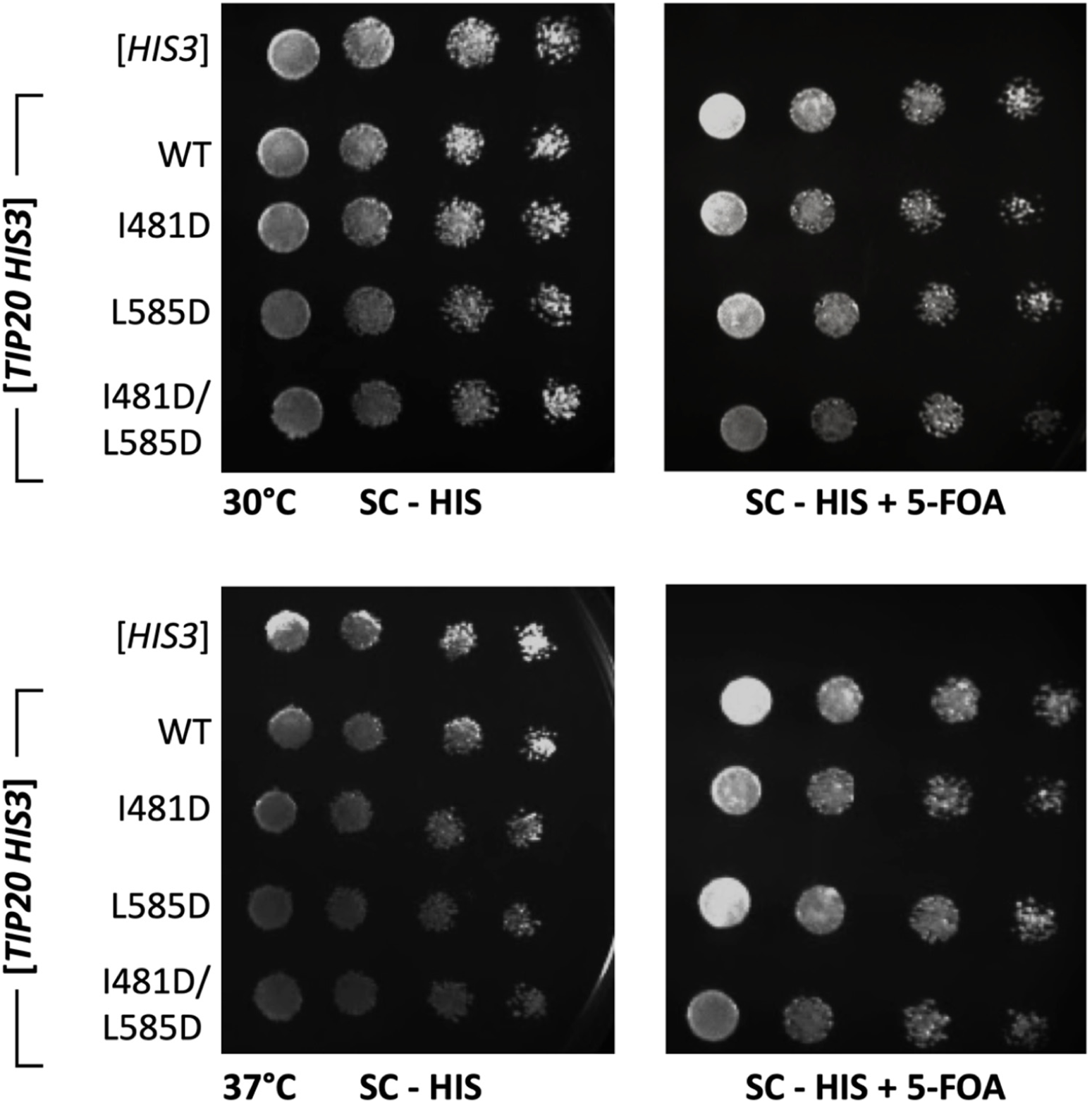
*S. cerevisiae* Tip20 interface mutants are viable. As in Figure 4, mutant alleles of *S. cerevisiae* Tip20, linked to His3, were tested for viability upon counterselection against a wild-type copy of Tip20, linked to Ura3. Although an empty His3 plasmid was not able to support viability, all alleles of Tip20 tested were viable, even at elevated temperature.

**Figure S6.**
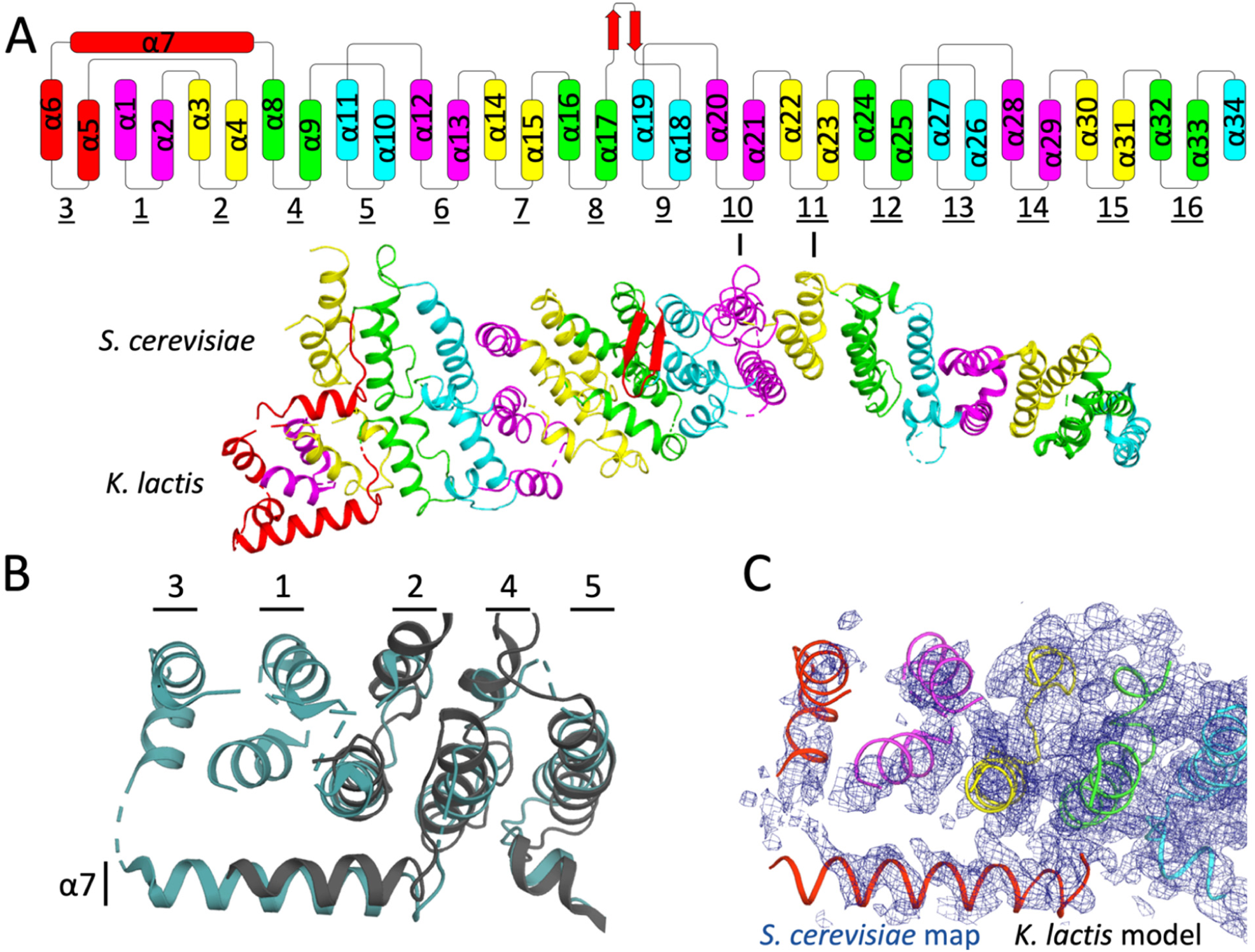
Additional N-terminal regions modeled for *K. lactis* Sec39. *A*, Above, topology of Sec39, with color-coding to illustrate the correspondence between individual α-helices and hairpins. Below, *S. cerevisiae* and *K. lactis* Sec39 superimpose with an RMSD of 3.7 Å over 436 residues. Superimposition at the C-terminus shows that the two structures differ primarily due to bending between layers 10 and 11 (indicated). *B*, Four additional helices were modeled at the N-terminus of *K. lactis* Sec39 (cyan), representing hairpins 1 and 3. In addition, the N-terminus of α7 was extended relative to *S. cerevisiae* Sec39 (3K8P, gray). The two structures are superimposed at the N-terminus. *C*, Contoured at 1 σ, the *S. cerevisiae* 2Fo-Fc map (3K8P, blue mesh) contains weak density, not previously interpreted, corresponding to hairpins 1 and 3 as modeled in the *K. lactis* structure.

**Figure S7.**
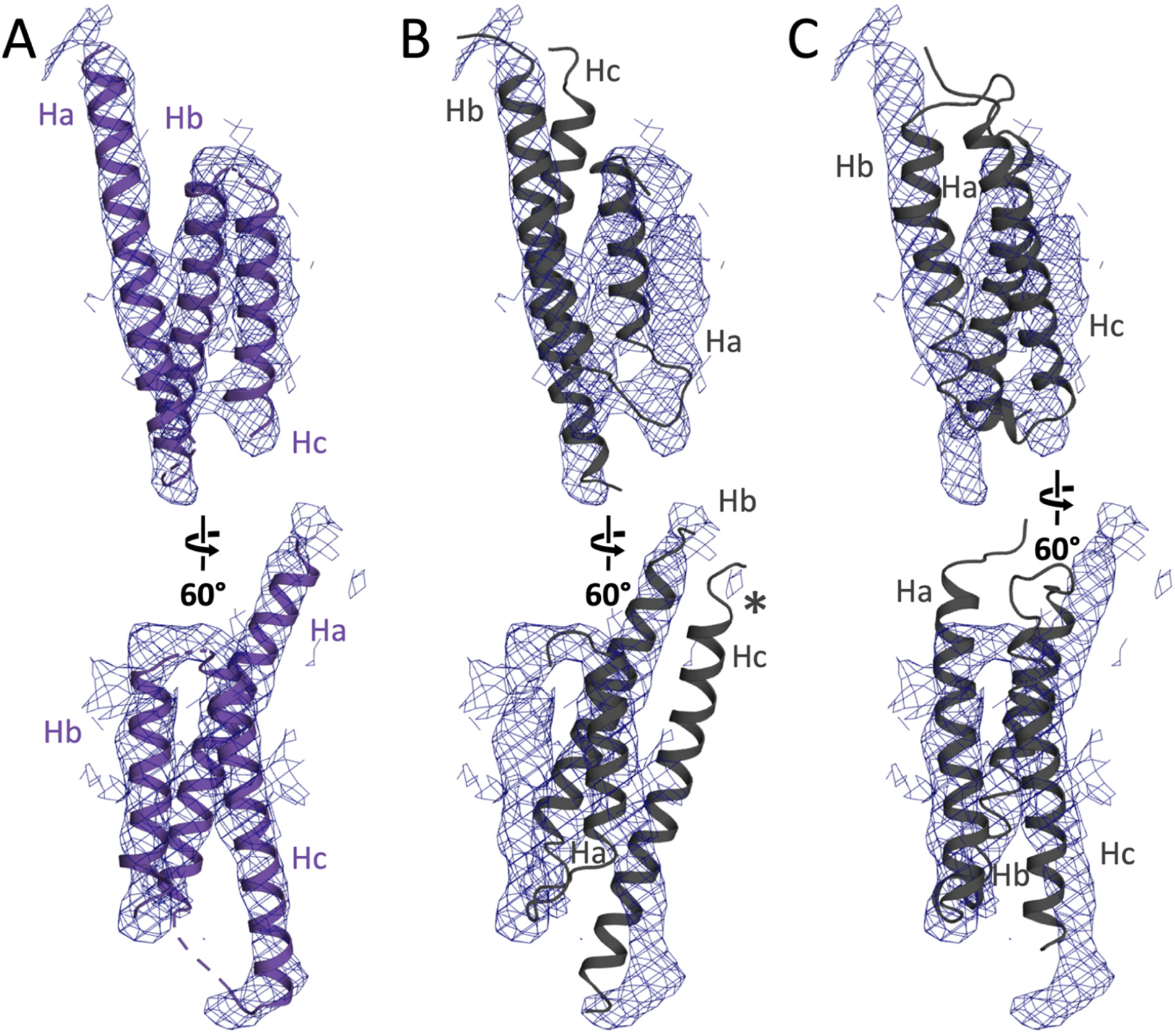
Electron density for *K. lactis* Use1 is consistent with an Habc domain topology. *A*, Contoured at 1 σ, the *K. lactis* 2Fo-Fc SNARE omit map (blue mesh) contains density for three α-helices, here modeled based on the top scoring molecular replacement solution, *S. cerevisiae* Vti1 (3ONJ, gray). *B*, An alternative molecular replacement solution, obtained by using *E. gossypii* Sec20 as a search model, failed to occupy all three helical densities. The starred helix instead clashes with hairpins 1 and 3 of Sec39. *C*, A second alternative molecular replacement solution, obtained by using *S. cerevisiae* Vam3 (1HS7) as a search model, failed to recapitulate inter-helix connectivities present in the electron density map. See also Table S3.

**Figure S8.**
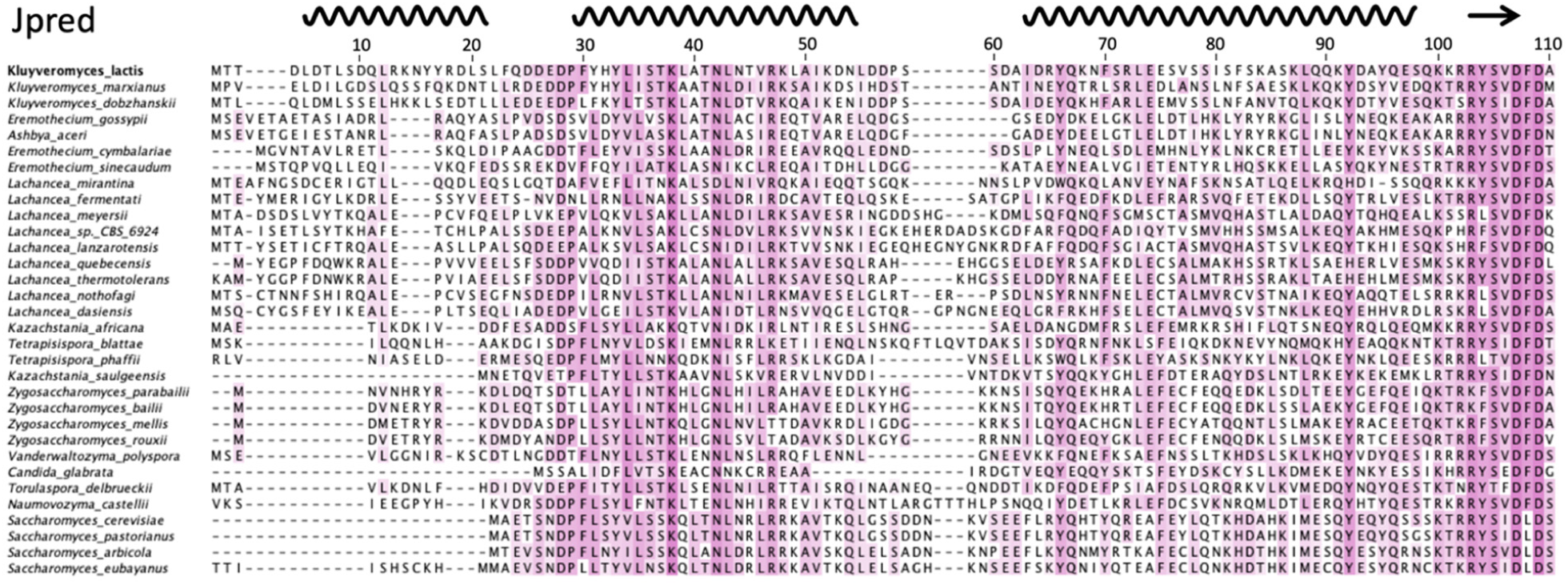
Secondary structure prediction suggests that the N-terminal domain of Use1 is α-helical. Secondary structure prediction (61) identified three putative α-helices in the N-terminal domain of *K. lactis* Use1. Because the first helix in the sequence is not conserved among some orthologues, including *S. cerevisiae*, it is possible that the long Ha helix observed in Use1 comprises the first two predicted helices. Sequence conservation is shown in purple.

**Table S1.**
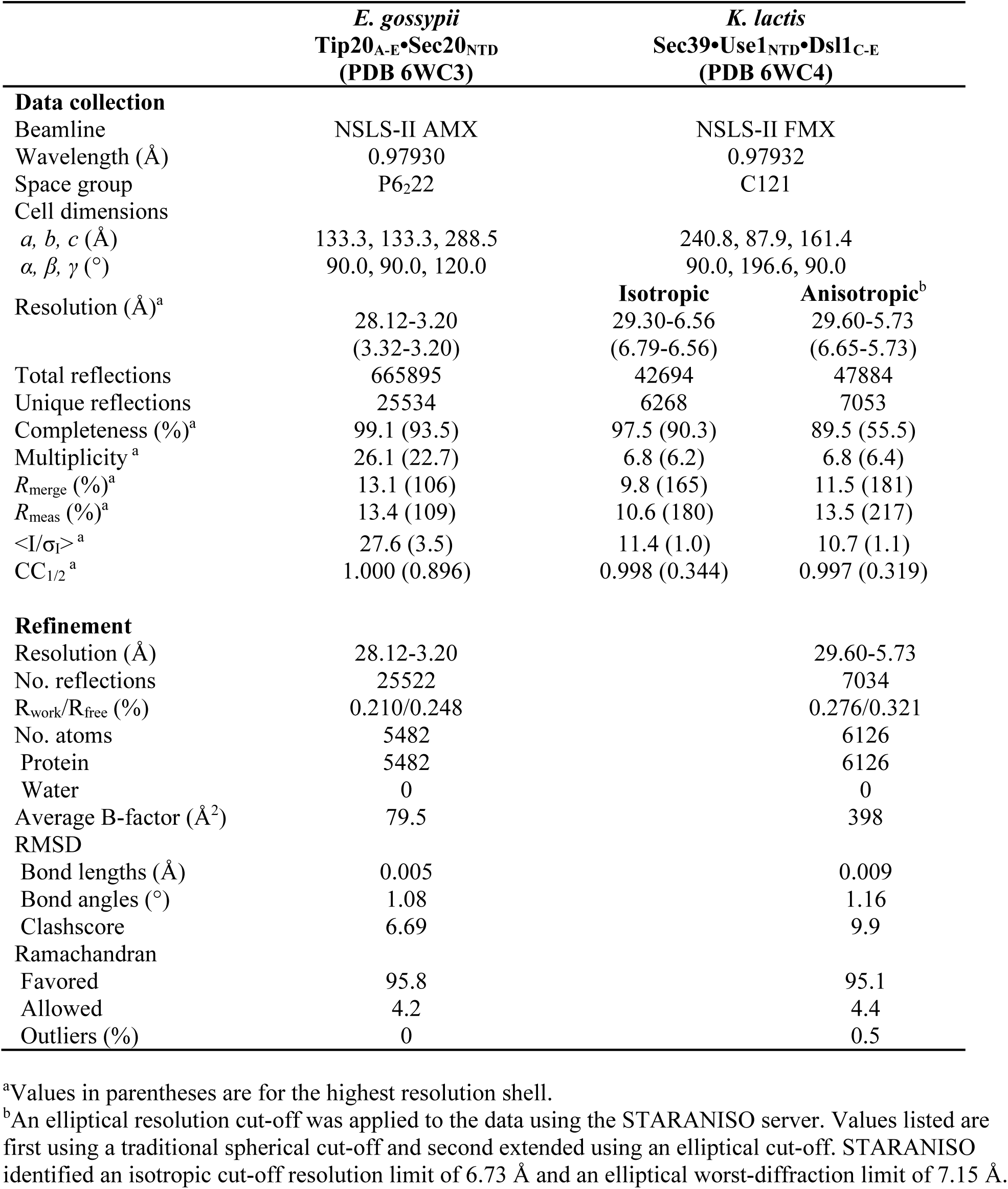
Summary of crystallographic parameters.

**Table S2.**
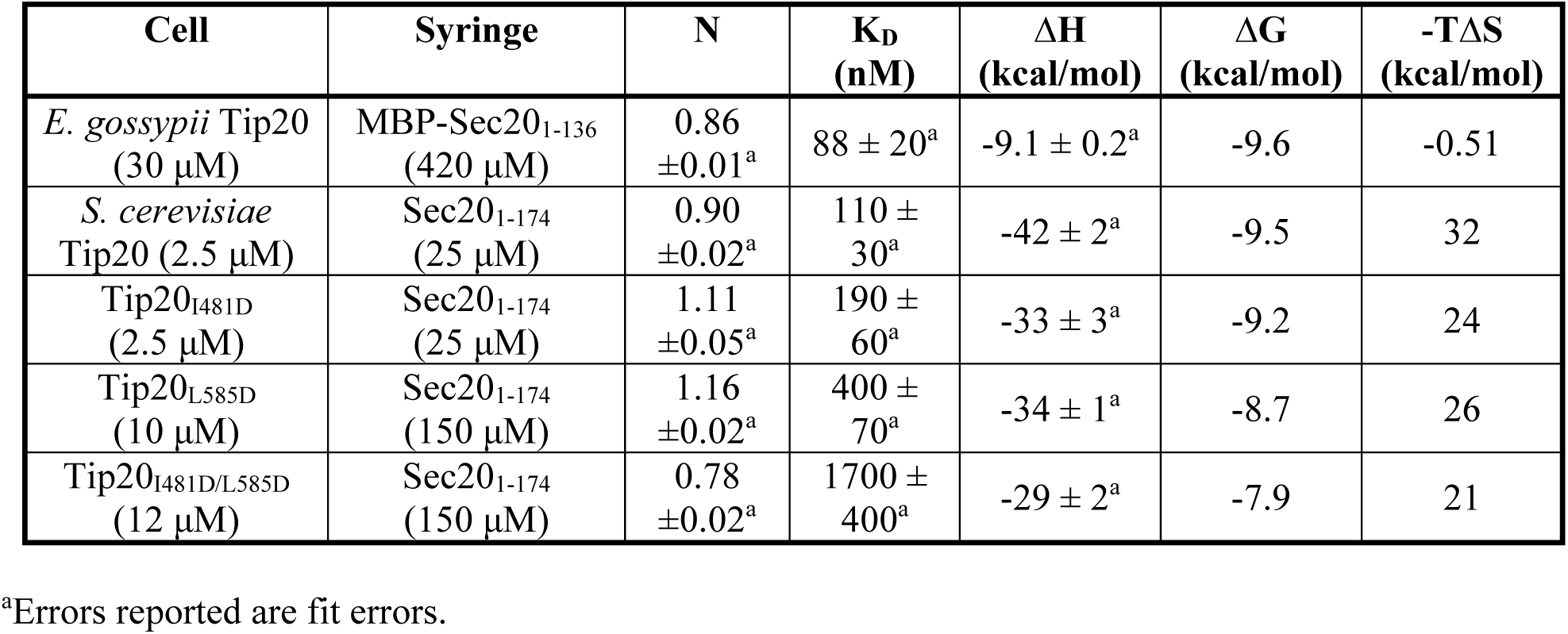
Isothermal titration calorimetry fit parameters of *S. cerevisiae* Tip20 mutants.

| Cell | Syringe | N | K<sub>D</sub><br>(nM) | ΔH<br>(kcal/mol) | ΔG<br>(kcal/mol) | -TΔS<br>(kcal/mol) |
| --- | --- | --- | --- | --- | --- | --- |
| *E. gossypii* Tip20<br>(30 μM) | MBP-Sec20<sub>1-136</sub><br>(420 μM) | 0.86<br>±0.01<sup>a</sup> | 88 ± 20<sup>a</sup> | -9.1 ± 0.2<sup>a</sup> | -9.6 | -0.51 |
| *S. cerevisiae*<br>Tip20 (2.5 μM) | Sec20<sub>1-174</sub><br>(25 μM) | 0.90<br>±0.02<sup>a</sup> | 110 ±<br>30<sup>a</sup> | -42 ± 2<sup>a</sup> | -9.5 | 32 |
| Tip20<sub>I481D</sub><br>(2.5 μM) | Sec20<sub>1-174</sub><br>(25 μM) | 1.11<br>±0.05<sup>a</sup> | 190 ±<br>60<sup>a</sup> | -33 ± 3<sup>a</sup> | -9.2 | 24 |
| Tip20<sub>L585D</sub><br>(10 μM) | Sec20<sub>1-174</sub><br>(150 μM) | 1.16<br>±0.02<sup>a</sup> | 400 ±<br>70<sup>a</sup> | -34 ± 1<sup>a</sup> | -8.7 | 26 |
| Tip20<sub>I481D/L585D</sub><br>(12 μM) | Sec20<sub>1-174</sub><br>(150 μM) | 0.78<br>±0.02<sup>a</sup> | 1700 ±<br>400<sup>a</sup> | -29 ± 2<sup>a</sup> | -7.9 | 21 |

<sup>a</sup>Errors reported are fit errors.

**Table S3.**
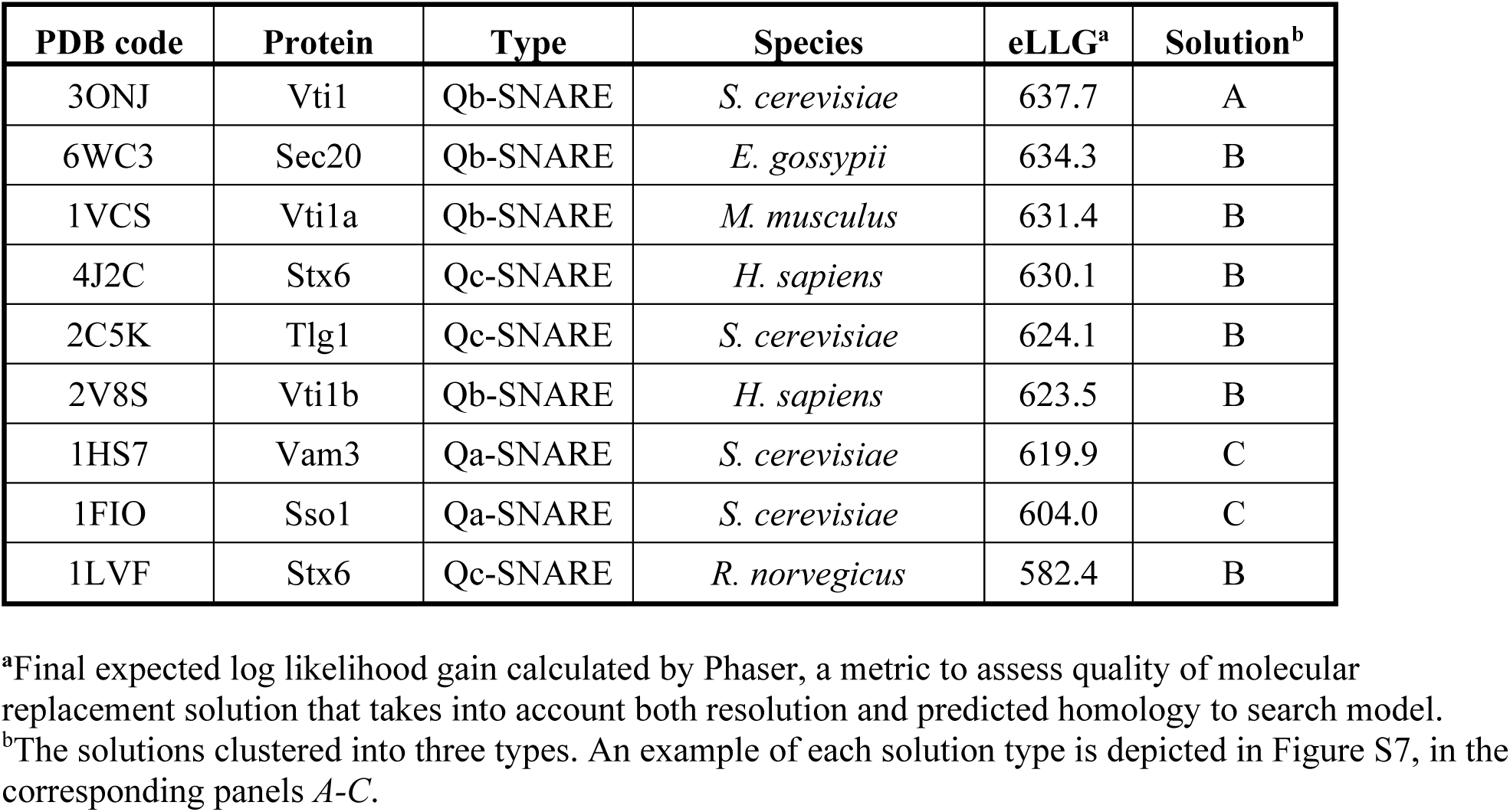
Alternative Use1 Habc domain molecular replacement solutions.

